# The evolution of splicing: transcriptome complexity and transcript distances implemented in *TranD*

**DOI:** 10.1101/2021.09.28.462251

**Authors:** Adalena Nanni, James Titus-McQuillan, Oleksandr Moskalenko, Francisco Pardo-Palacios, Zihao Liu, Ana Conesa, Rebekah L. Rogers, Lauren M McIntyre

**Author notes:** Corresponding author: Lauren McIntyre, Rebekah L. Rogers. First author (Adalena Nanni, James Titus-McQuillan).

## Abstract

Alternative splicing contributes to organismal complexity. Comparing transcripts between and within species is an important first step toward understanding questions about how evolution of transcript structure changes between species and contributes to sub-functionalization. These questions are confounded with issues of data quality and availability. The recent explosion of affordable long read sequencing of mRNA has considerably widened the ability to study transcriptional variation in non-model species. In this work, we develop a computational framework that uses nucleotide resolution distance metrics to compare transcript models for structural phenotypes: total transcript length, intron retention, donor/acceptor site variation, alternative exon cassettes, alternative 5’/3’ UTRs are each scored qualitatively and quantitatively in terms of number of nucleotides. For a single annotation file, all differences among transcripts within a gene are summarized and transcriptome-level complexity metrics: number of variable nucleotides, unique exons per gene, exons per transcript, and transcripts per gene are calculated. To compare two transcriptomes on the same co-ordinates, a weighted total distance between pairs of transcripts for the same gene is calculated. The weight function proposed has larger penalties for intron retention and exon skipping than alternative donor/acceptor sites. Minimum distances can be used to identify both transcript pairs and transcripts missing structural elements in either of the two annotations. This enables a broad range of functionality from comparing sister species to comparing different methods of building and summarizing transcriptomes. Importantly, the philosophy here is to output metrics, enabling others to explore the nucleotide-level distance metrics. Single transcriptome annotation summaries and pairwise comparisons are implemented in a new tool, *TranD,* distributed as a *PyPi* package and in the open-source web-based Galaxy (www.galaxyproject.org) platform.

## Introduction

Many eukaryotic genes are transcribed into multiple mRNAs through alternative splicing (AS) leading to differing functional properties (e.g., Roretz and Gallouzi 2008) and/or encoded proteins from the same gene (e.g., Gilbert 1978; Gilbert, et al. 1997; Vibranovski, et al. 2005; Frankish, et al. 2012). This differential processing has been hypothesized to be a primary mechanism of protein diversity in higher eukaryotes (e.g., Gilbert 1978; Hirai, et al. 2004; McGuire, et al. 2008; Lu, et al. 2010; Mudge, et al. 2011), and there is a relationship between organismal complexity and splicing (Chen, et al. 2014). Dramatic differences in splicing events among tissues, environments, cell types and developmental stages within a single species are well documented across the tree of life (e.g., Akam and Martinezarias 1985; Bell, et al. 1988; Bermingham and Scott 1988; O’Connor, et al. 1988; Graveley 2001; Celniker, et al. 2009; Levin, et al. 2012; Klepikova, et al. 2016; Newman, et al. 2017; Xiong, et al. 2018; Aguet, et al. 2020). Yet, whether alternative splicing is conserved has conflicting evidence with suggestions of both rapid divergence (Roy and Irimia 2009; Barbosa-Morais, et al. 2012; Merkin, et al. 2012) and conservation in splicing regulation. Specifically, sex determination genes in insects with sex specific splicing (Burtis and Baker 1989; Telonis-Scott, et al. 2009; Salz and Erickson 2010; Graveley, et al. 2011; Rogers, et al. 2021) and developmental stage specific splicing (Irimia, et al. 2009; Gao, et al. 2021; Mazin, et al. 2021).

One potential explanation for the apparent contradiction in the observations of conservation and divergence in splicing, is in the evolution of the spliceosome itself. Multiple different molecular mechanisms contribute to the process (e.g., Tolstrup, et al. 1997; Lorkovic and Barta 2002; Zhu, et al. 2003). The spliceosome is a large complex with some conserved splicing factors (Kreivi and Lamond 1996; Collins and Penny 2005; Jangi and Sharp 2014) and a plethora of context specific factors (Jangi and Sharp 2014), some of which may be shared along specific evolutionary branches (McManus, et al. 2014). For example, sex-determination (Boggs, et al. 1987), smooth muscle MBNL cascade, T-cell states, neuronal Nova splicing regulation, and brains of primates (Mazin, et al. 2018). Further, observations of conservation of sex (e.g., Dauwalder, et al. 1996) and neurological (e.g. Srivastava, et al. 2021) splicing factors between human and fly indicate that there are potentially more conserved regulatory mechanisms of splicing, even as genes that are regulated by these factors will almost certainly differ depending on the larger transcriptional regulatory environment.

There is growing interest in how to approach transcripts in a phylogenetic context (Zea, et al. 2021). Modes of splicing variation may differ across taxa (Clark and Thanaraj 2002; Kan, et al. 2002; Ner-Gaon, et al. 2004; Mei, et al. 2017; Freese, et al. 2019; Jia, et al. 2020). For example, more cassette exons are conserved in nematodes (Irimia, et al. 2008) than mammals (Modrek and Lee 2003; Pan, et al. 2005; Nurtdinov, et al. 2007), where the elapsed amount of evolutionary time is in approximately the same range (Waterston, et al. 2002; Stein, et al. 2003; Cutter 2008). Cassette exons rely on the concept of genome conservation (Irimia, et al. 2009) and exon skipping rates vary across eukaryotes (e.g., Grau-Bove, et al. 2018). Alternative donor and acceptor sites have been suggested to be subject to more rapid evolution than exon cassettes as they are part of an evolutionary process on the intron/exon boundaries (e.g., Mazin, et al. 2018). There is more use of intron retention in plants compared to mammals (see Martin, et al. 2021). As the number of annotated genomes grows there is an opportunity to clarify how well splicing mechanisms are conserved and what exceptions may be present in newly sequenced taxa (Irimia, et al. 2009).

Short reads identify local splicing changes, rather than whole transcripts. While there are a plethora of approaches to transcript reconstruction from short reads, there is consensus that these approaches have limitations in the ability to accurately reconstruct and quantify transcripts (Steijger, et al. 2013; Freedman, et al. 2021) especially in lower coverage sequence data (MaManes 2018). Orthologous exons (Clark, et al. 2007; Fu and Lin 2012; Douzery, et al. 2014), junction event orthology (Sheth, et al. 2006; Chorev, et al. 2016; Mei, et al. 2017), and estimates of percent spliced in (Katz, et al. 2010) provide local comparisons. As long-read technology error rates are decreasing and costs are plummeting (Amarasinghe, et al. 2020), the opportunities for experimentally assaying whole transcripts are an exciting opportunity for a diverse community of scientists. Existing tools combine annotations and make non-redundant selections (Trapnell, et al. 2012), blend different types of data for transcript models, and increasingly estimate transcriptomes from long reads (Tang, et al. 2020). What is missing is a way to make explicit comparisons among these approaches.

Here, we introduce *TranD,* an open source *PyPi* package (https://github.com/McIntyre-Lab/TranD), also available as a wrapped tool in Galaxy (www.Galaxyproject.org), that identifies and quantitates nucleotide-level structural differences. For a single annotation file (1GTF) genelevel metrics include intron retention, exon skipping, alternative donor/acceptor sites, and 5’/3’ UTR variation. For each gene, the numbers and percentages of variable nucleotides, exons, and transcripts are reported. A transcriptome summary of the distribution of exons per transcript; transcripts per gene; and exons per gene is complemented by visualizations of the metrics. In addition to gene-level metrics, all pairs of transcripts within a gene can be compared, facilitating identification of transcripts that share specific structural phenotypes. In addition to descriptions of structural phenotypes within and across genes for a single annotation, metrics that compare two annotations (2GTF) have been implemented. These metrics require both annotations to be on the same set of genome coordinates. When comparing a pair of transcripts, all structural phenotypes (number of alternate donor/acceptor sites, intron retentions, alternative exons cassettes, alternative 5’ and 3’ ends) and the shared and unshared nucleotides are reported. A minimum pair is identified using a sequential process where first intron retentions and exon skipping are minimized, then donor and acceptor variation and finally 5’ and 3’ variation.

These metrics are all individually reported enabling a wide range of researchers to quickly identify pairs of similar and divergent transcripts. The possible applications have a wide breadth of utility. Here we present examples to illustrate the utility of *TranD:* metrics for the *D. melanogaster, Z. mays,* and *C. elegans* genomes illustrate the 1GTF gene and pairwise modes; the refinement of *D. yakuba* transcriptome annotation and a comparison of two methods for estimating transcriptomes from long reads, Isoseq3 (https://github.com/PacificBiosciences/IsoSeq) and FLAIR (Tang, et al. 2020), illustrate the 2GTF pairwise mode.

## New approaches

### Variation in structural phenotypes among genes (TranD 1GTF gene-mode Figure 1A)

**Figure 1.**
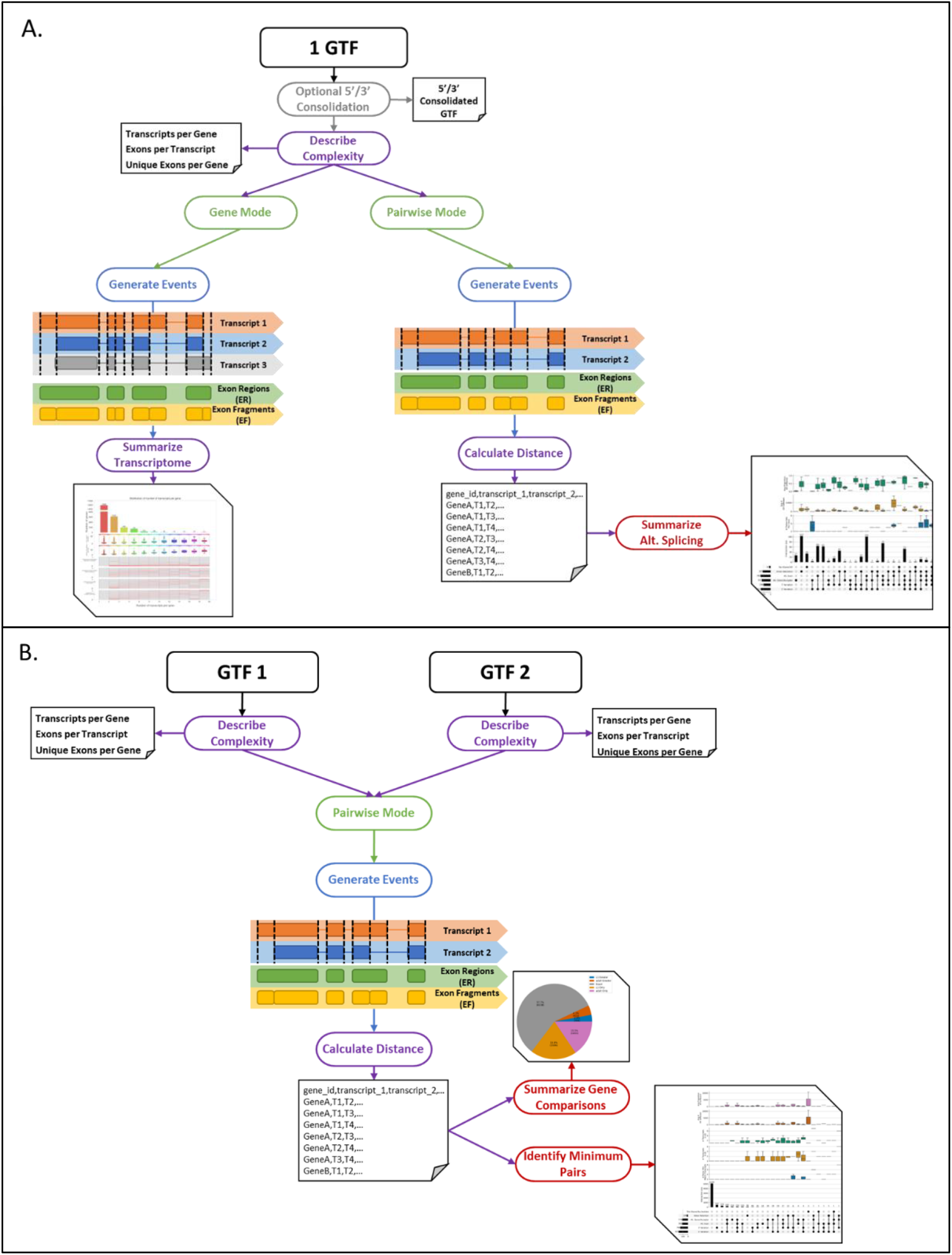
Transcript Distance *(TranD)* can be used to describe and quantify structural differences within a single transcriptome (A) or between two transcriptomes (B).

For genes with more than 1 transcript, structural phenotypes are enumerated using the approach described in Newman et. al. (Newman, et al. 2018). The union of all transcripts is used to define the transcript space. The transcript space is then annotated for variation in the following structural phenotypes: intron retentions, donor/acceptor sites, alternative exons, and alternative 5’/3’ ends. The transcript space is defined by the 5’ most start, and 3’most end. Here we are focused just on the nucleotide identity in the transcripts and not functional annotation. For gene g, the total number of nucleotides in the transcript space is N_g_. The number of nucleotides that vary is V_g_, and the percentage of variable nucleotides is 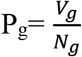. The distributions of P_g_ across the genome provides insight on the variability of the transcriptome and can be compared to other transcriptomes to assess the impact of splicing in a particular species.

### Variation in structural phenotypes among transcripts (TranD 1GTF pairwise-mode Figure 1A)

Between a pair of transcripts T_j_ and 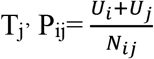 (percent identity) is a metric of the difference in the number of nucleotides between the two transcripts, while 1-P_ij_ is the proportion of the transcript space shared. Additionally, the presence/absence of nucleotides is tracked for each transcript. For example, which transcript in the pair contains the intron retention. These metrics are calculated for all possible pairs of transcripts within a gene (Supplemental Figure 1). Then for each pair across the genome the unique combination of structural phenotypes are summarized and the distribution of the nucleotide variation for pairs with the same set of structural phenotypes are displayed.

### Distance between transcripts in two annotations (2GTF, all pairwise Figure 1B)

To quantify nucleotide level variation between two annotations, the transcriptomes must be on the same co-ordinate system and have identical gene names. For each gene g, transcript j, from GTF1 is compared to transcripts 1 to k in annotation GTF2; for all j in gene g. For all pair-wise comparisons between transcripts of gene g in GTF1 and GTF2, there is one pair that has a minimum distance. As we expect a higher proportion of nucleotides to differ in intron retentions compared to alternative/donor and acceptors we do not use Pkj to select the minimum distance among pairs. Instead, we use a sequential procedure that first compares exon regions, then junctions and finally P_kj_. We do not assume the min_TD_k,j_ to be the same as the min_TD_j,k_. The minimum distance is indicated a binary 0/1 variable (*flag_minimum_match_[A]*), where A represents the name given to each annotation (“d1” and “d2” by default). When the minimum distance for transcript j is transcript k and the minimum distance for transcript k is transcript j, the transcripts are a reciprocal minimum pair *(flag_recip_min_malch* = 1) (Supplemental Figure 2). We do not argue that this is the best minimum distance metric, indeed this remains to be proven. As reasonable scientists may disagree about this order of priority, all individual metrics and nucleotide variability are output enabling other versions of the minimum distance between two transcripts min_TD_kj_ to be readily calculated (Figure 2). The number of genes and transcripts that are exclusive to one annotation are tracked and recorded. When for the same gene, there are more transcripts for one of the annotations, there is the possibility that the annotation with the smaller number is a complete subset of the larger number of transcripts. The definition of the reciprocal pair does not imply that the distance between the two transcripts is small.

### Complexity metrics

We propose three general metrics of complexity that do not rely on comparing transcripts directly, but rather describe the global structural phenotypes of transcriptomes: 1) the number of transcripts per gene (TpG); 2) the number of exons per transcript (EpT); and 3) the number of unique exons (exons with unique genomic coordinates) per gene (EpG). From each these metrics, distributions across genes can be examined. Summary statistics such as mean, median, variance, inter-quartile range are calculated to provide insights into the shapes of these distributions. These metrics are calculated for each transcriptome regardless of which scenario is being considered and can be used to compare transcriptomes on different coordinate systems.

## Results and discussion

### Structural phenotypes provide a useful summary of the mechanisms of splicing

In *D. melanogaster* (FlyBase r6.17, http://ftp.flybase.net/releases/FB2017_04/dmel__r6.17/gtf/dmel-all-r6.17.gtf.gz) there are 17,479 genes and most (58%) are single transcript genes. Of the 7,386 genes with more than 1 transcript ~16% are variable only at the 3’ end and 23% have a single type of structural variation (Supplemental Figure 3), indicating that the majority of genes with alternative splicing show multiple differences in structural phenotypes. The structural phenotypes contributing to the overall nucleotide variability are a balance between the different types of structural variation and overall the majority of genes have low proportions of nucleotide variability (less than 20%) (Figure 3A). Taking a closer look at the data by focusing on all possible pairs of transcripts (Figure 3B) we see 321 pairs (from 30 genes, Supplemental Figure 3) share no nucleotides in common. These genes may have longer isoforms combining exons from each member of the pair, even when the individual transcripts of the pair do not overlap. Hence, they are annotated as part of a single gene. 30% of transcript pairs have a single structural difference between them while 70% have more than 1 structural difference with the most frequent (60% of all pairs) being alternative exons coupled with 5’ and/or 3’ variation (Figure 3B).

**Figure 2.**
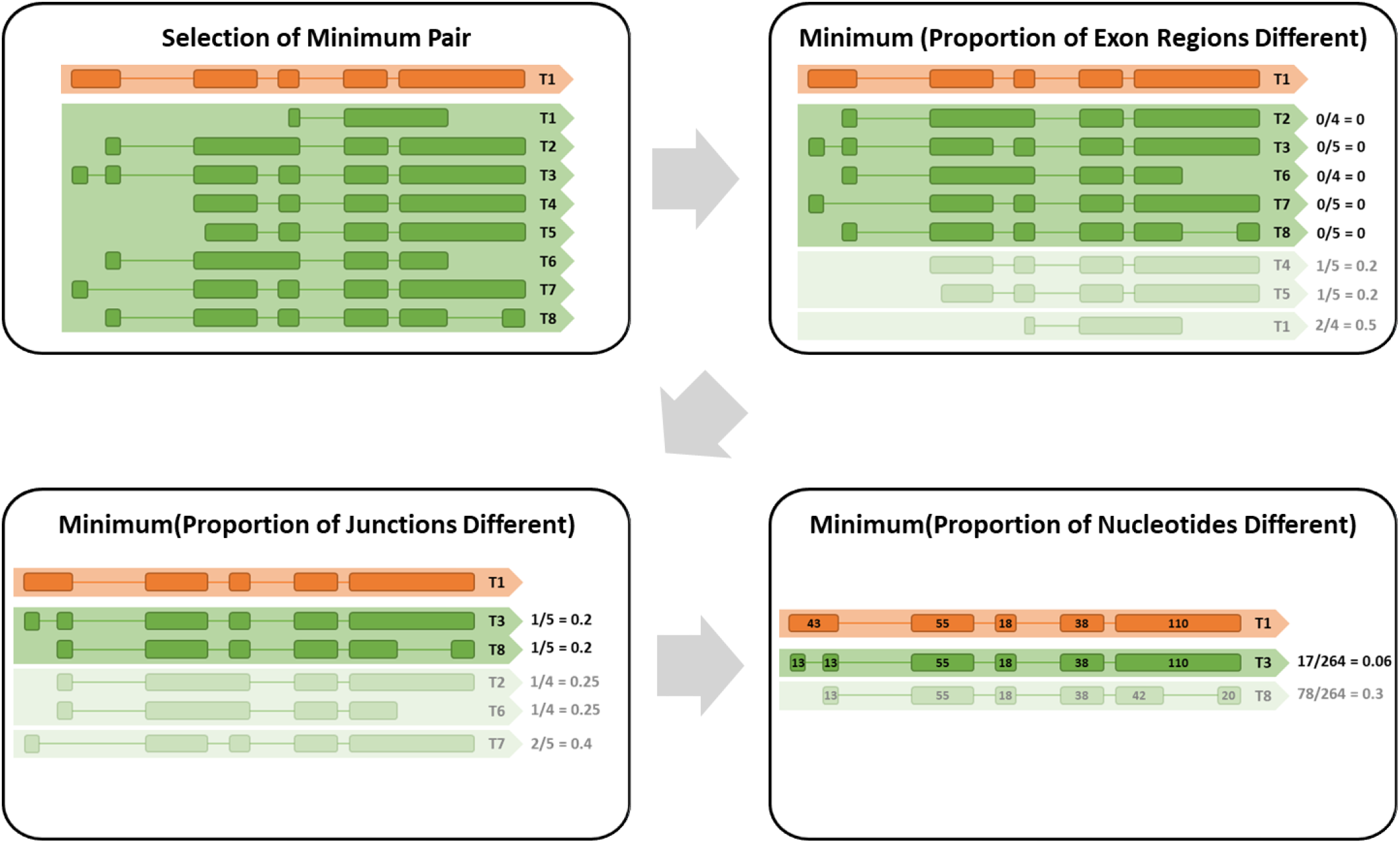
When comparing two annotations, distances between all possible pairs of transcripts are calculated. A minimum pair is determined in a sequential ranking of minimum distances for each distance in this order: the proportion of exon regions different, proportion of junctions different, and proportion of nucleotides different. Each panel, starting with the top left, demonstrates the computation of the individual distance and selection of the minimum. If more than one transcript is determined to be the minimum after all metrics are calculated, transcript names are sorted and the first is selected.

**Figure 3.**
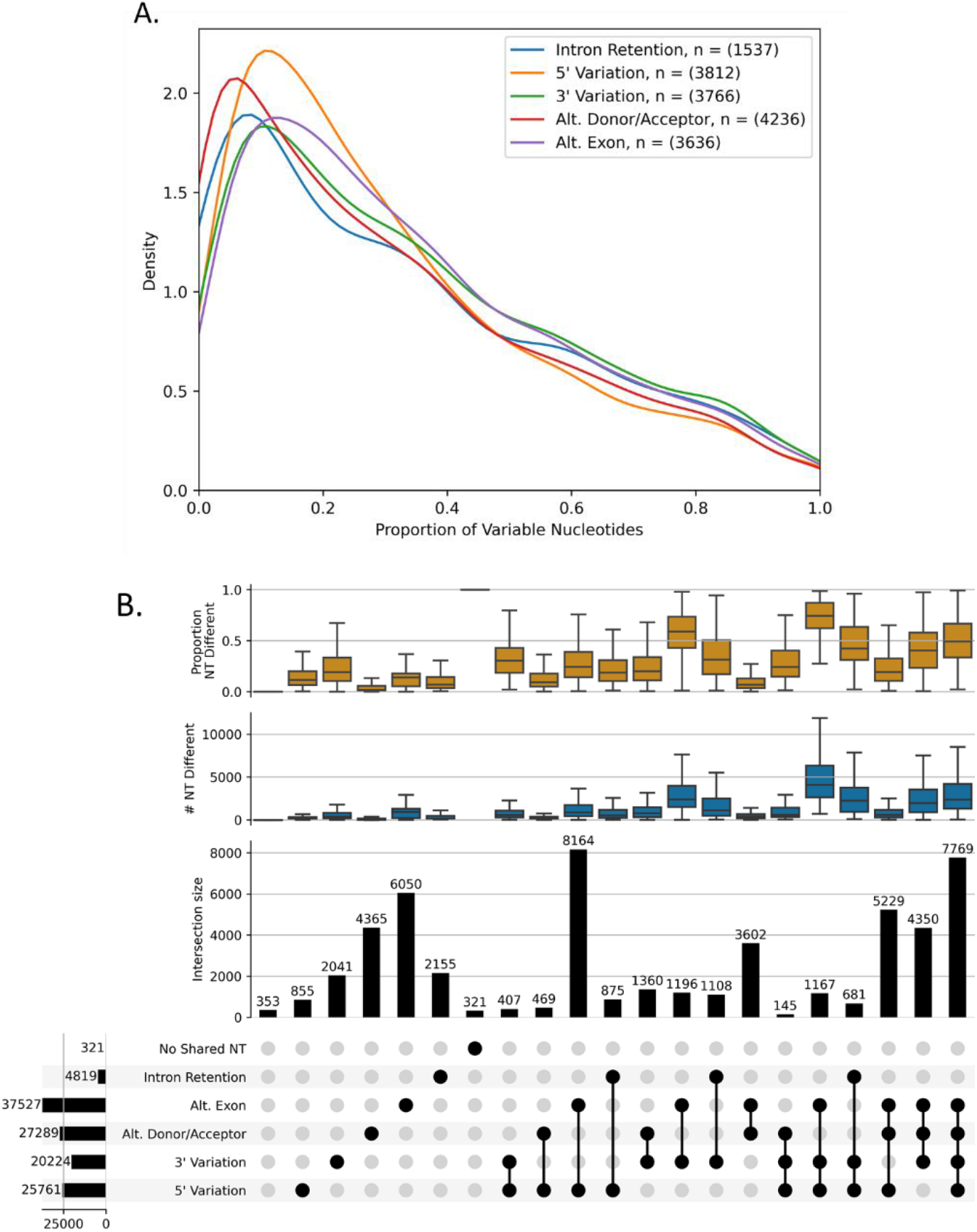
Structural phenotypes of *D. melanogaster* FlyBase 6.17 reference transcriptome. (A) Distributions of nucleotide variability across genes that contain each of the indicated alternative splicing categories. Genes that contain more than one category will be included in more than one distribution. (B) Number of transcript pairs with the specified types of alternative splicing indicated by the black dots below the histogram of transcript pair counts (n = 52662 transcript pairs, in 7386 multi-transcript genes). Box plots represent the number (blue) and proportion (orange) of nucleotides different between the pair. Transcript pairs with “No Shared NT” have nonoverlapping coordinates.

### Structural phenotypes differ between species

We compare the transcriptomes of classic model species *D. melanogaster* (FlyBase r6.17, Supplementary Figure 4), *C. elegans* (C. elegans Sequencing Consortium 1998) (WBcel235 annotation, Supplementary Figure 5) and a model plant known for its genome complexity, *Z. Mays* (Yang, et al. 2017) (Mo17 YAN annotation, Supplementary Figure 6) and H. sapiens (RefSeq v38, Supplementary Figure 7)*. D. melanogaster,* and *C. elegans* annotations have more than 50% of the genes with only one transcript. *H. sapiens* and *Z*. *mays* have a larger proportion of genes with four or more transcripts. *Z. mays* has more annotated intron retentions, and these contribute substantially to variation in genes with larger transcript numbers. When there are 8 or more transcripts per gene, over 50% of all *Z. mays* transcripts contain an IR event, consistent with previous literature describing IR as the most prevalent class of alternative splicing in plants (reviewed in Barbazuk, et al. 2008; Marquez, et al. 2012). When we focus on the non-intron retention structural phenotypes, we find that *Z. mays* has proportionally more alternative donor/acceptor sites than *C. elegans.* However, *C. elegans* has proportionally higher numbers of alternative exons than *Z. mays*. *D. melanogaster* and *H. sapiens* are a balance with both alternate donor and acceptors and alternative exons cassettes. Our results suggest that complexity in *Z. mays* is driven by changes in splicing junctions, while *C. elegans* isoform variability may be driven by combinatorics of frequently used exons. Quantitative differences in transcripts increase as the number of transcripts increases in all four species.

Our results are concordant with prior research on spliceosomal differences and differential splicing in plants and animals. Plants are known to have high intron-retention within their spliceosome (Ner-Gaon, et al. 2004; Mei, et al. 2017; Freese, et al. 2019; Jia, et al. 2020). In humans only 2-5% of introns are retained during splicing across the genomes (Clark and Thanaraj 2002; Kan, et al. 2002). Plants have less well-defined borders between introns (Zhu, et al. 2003) and intron branch points (Tolstrup, et al. 1997). There are also more complex families of spliceosomal-associated serine arginine (SR) factors are possessed by plant lineages (Lorkovic and Barta 2002). Splicing signals vary between eukaryotic lineages where plants have higher non-GT/AG in plants than in animals (Ye, et al. 2017), possibly influencing alternate donor and acceptor sites.

### Comparing transcriptome annotations in D. Yakuba

In emerging genetic systems, developing annotation is a critical component to downstream analyses in molecular and evolutionary genetics. Uncertainty regarding annotation can complicate progress even with technological improvements of long molecule sequencing. Comparing transcripts precisely using distance metrics makes the job of iterating annotations in developing systems a fraction of the effort compared to the current state of the art. We demonstrate this with a comparison of annotation versions for *D. yakuba.* This case study provides a roadmap for implementing distance measures and transcriptome comparison metrics quickly and easily in *TranD* to compare annotation strategies in the rapidly growing field of non-model organism genetics.

Two annotations from the fruit fly species *D. yakuba* are compared: FlyBase r1.05 *D. yakuba* and RNA-seq based *D. yakuba* annotations (here on referred to a dyak-RR-revised) (Rogers, et al. 2014) using *TranD*. These two annotations were produced with different strategies comparative genomics (Clark, et al. 2007) and data based (Rogers, et al. 2014). These approaches are expected to capture different features of transcriptomes. Our goal was to leverage each of these methods to produce one more comprehensive annotation for *D. yakuba.* Using distance metrics, we compare **all pairs** of transcripts for each gene between the two annotations using *TranD*, a matrix of differences is output, as are the set of visualizations that compare the two annotations.

Between FlyBase r1.05 *D. yakuba* and dyak-RR-revised, the FlyBase annotations have more genes annotated and more transcripts while for genes present in dyak-RR-revised are more annotated exons per gene and per transcript. Across all 21237 genes in either annotation, only 8162 are identified in both annotations, 7907 genes are found only in FlyBase r1.05 and 5168 genes only in dyak-RR-revised (Figure 4). We augmented the complete FlyBase r1.05 annotation set with 25,022 transcripts across 16,069 genes with 3,102 transcripts corresponding to 3,089 novel gene sequences from the dyak_RR annotations. 3,911 transcripts from 2,875 genes with novel intron retentions or exon regions from the dyak_RR annotations were added to FlyBase r1.05 annotation. A total of 134 of these transcripts had at least one other transcript in dyak_RR annotations with only 5’ and 3’ variation. In these cases, the longer of the two transcripts was included. The FB105+ annotation file has a total of 32,035 transcripts and 19,158 putative genes. FB105+ annotation encompasses more complete information from past and current update. A similar operation could have been performed in cuff compare/cuff merge (Trapnell, et al. 2012), joining transcripts based on overlapping exons or genomic locations. However, by inferring structural phenotypes that differentiate transcripts, we can identify transcripts that add information about exon usage in the reference. FB105+ offers greater precision on isoform prediction than either dataset alone and retains information that may otherwise be lost in compression.

**Figure 4.**
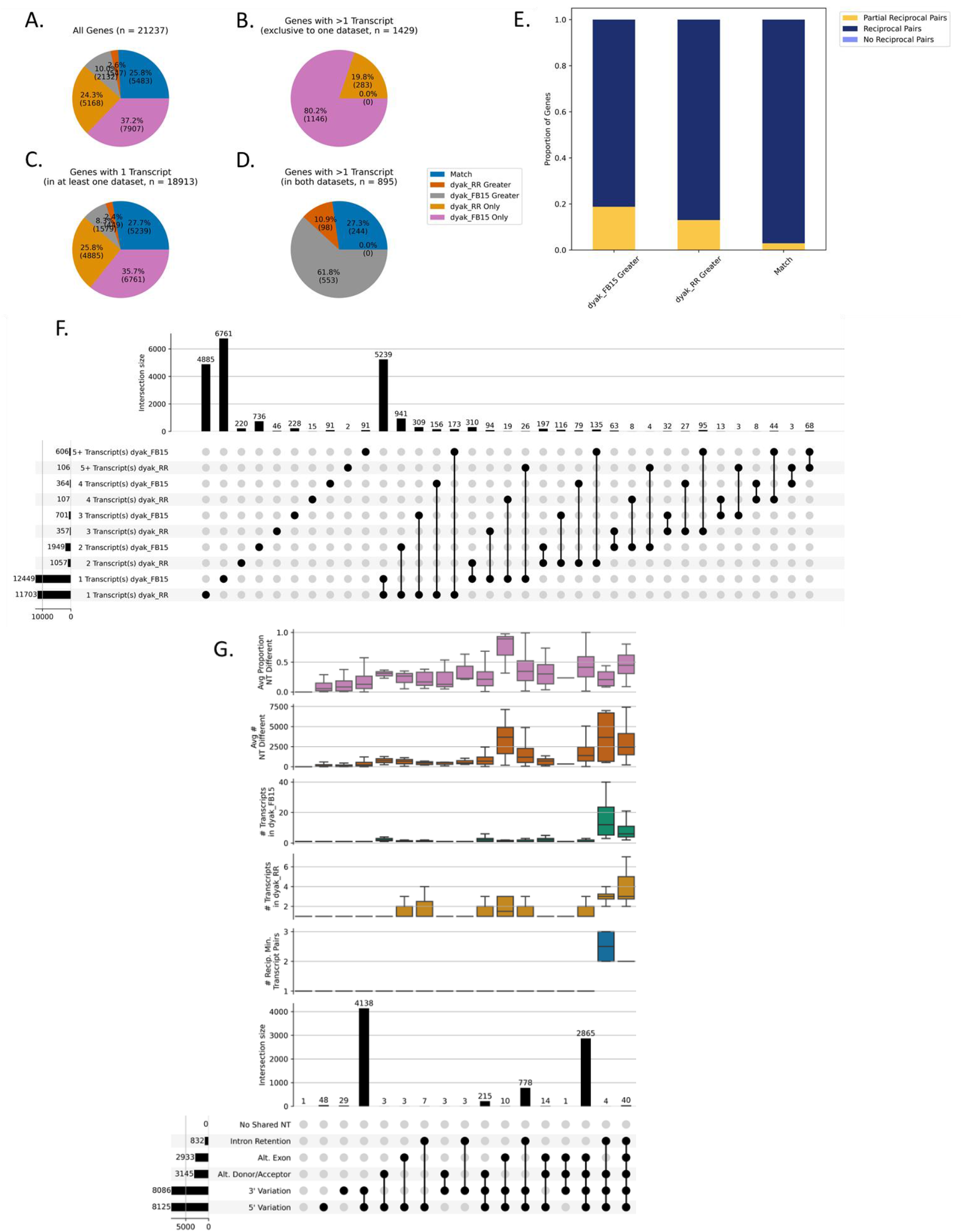
Transcript models from *D. yakuba* Rogers, et al. 2014 annotation (here on referred to as dyak-RR-revised) and *D. yakuba* FlyBase r1.05 (here on referred to as dyak-FB105) are compared using *TranD.* (A-D) Percent of genes with equal number of transcripts between dyak-RR-revised and dyak-FB105 (Match, blue), more transcripts in dyak-RR-revised compared to dyak-FB105 (dyak-RR-revised Greater, dark orange), more transcripts in dyak-FB105 compared to dyak-RR-revised (dyak-FB105 Greater, gray), and genes exclusive to dyak-RR-revised (dyak-RR-revised Only, light orange) or dyak-FB105 (dyak-FB105 Only, pink). (E) Within genes with equal numbers of transcripts in dyak-RR-revised and dyak-FB105 (Match), the number of genes where all transcripts have reciprocal minimum matches (Match: Reciprocal Pairs, n = 5323), at least one but not all transcript pairs are reciprocal minimum matches (Match: Partial Reciprocal Pairs, n = 160), and no pairs are reciprocal minimum matches (Match: No Reciprocal Pairs, n = 0). Within genes with more transcripts in dyak-RR-revised compared to dyak-FB105 (dyak-RR-revised Greater), the number of genes where all transcripts in dyak-FB105 have reciprocal minimum matches to a subset of dyak-RR-revised (dyak-RR-revised Greater: Reciprocal Pairs, n = 476), at least one but not all transcript pairs are reciprocal minimum matches (dyak-RR-revised Greater: Partial Reciprocal Pairs, n = 71), and no pairs are reciprocal minimum matches (dyak-RR-revised Greater: No Reciprocal Pairs, n = 0). Within genes with more transcripts in dyak-FB105 compared to dyak-RR-revised (dyak-FB105 Greater), the number of genes where all transcripts in dyak-RR-revised have reciprocal minimum matches to a subset of dyak-FB105 (dyak-FB105 Greater: Reciprocal Pairs, n = 1732), at least one but not all transcript pairs are reciprocal minimum matches (dyak-FB105 Greater: Partial Reciprocal Pairs, n = 400), and no pairs are reciprocal minimum matches (dyak-FB105 Greater: No Reciprocal Pairs, n = 0). (F) Rows are the number of transcripts in dyak-RR-revised and dyak-FB105. Columns indicate the genes with the unique combination of transcript numbers in the two annotations. Columns with only a single black dot occur for genes present only in dyak-RR-revised (n = 5168) or dyak-FB105 (n = 7907). (G) For genes present in both annotations and at least 1 reciprocal minimum pair the differences in transcripts in reciprocal minimum pairs of the dyak-RR-revised and dyak-FB105 (n = 8162 genes with 8634 reciprocal minimum pairs). The Y axes are counts of structural variation for each pair; distributions of the number of reciprocal minimum pairs in each gene (blue); number of transcripts in dyak-RR-revised (orange) and dyak-FB105 (green), and the average number (brown) and proportion (purple) of nucleotides different between the pairs. Genes with “No Shared NT” have a pair of transcripts with nonoverlapping coordinates.

## Annotations depend upon data quality

### Data processing pipelines affect annotation

Long read transcriptomic data are now affordable and can used as an unbiased look at the transcriptome. However, these technologies are not without error, and some technical obstacles still need to be overcome in using these data to estimate transcripts. A 2020 review identified 36 analysis tools for processing long read RNA sequencing and estimating transcripts (Amarasinghe, et al. 2020). The most commonly used include the PacBio IsoSeq3 cluster function (https://github.com/PacificBiosciences/IsoSeq) which does not use a reference and FLAIR cluster (Tang, et al. 2020) initially developed for Oxford Nanopore long reads which leverages a reference annotation.

We used *TranD* 2GTF all pairwise to compare transcripts from the GTF files of annotated transcriptomes estimated from these two tools from the same long read data. For this particular comparison we used 4 previously published datasets in *C. elegans* (L1) (Roach, et al. 2020) and *Z. mays* (root) (Wang, et al. 2020). We found that IsoSeq3 (which does not use a reference) returned more transcripts per gene than FLAIR (which uses a reference) from the same set of long reads. A higher degree of 5’ and 3’ variation was also observed in IsoSeq3 references compared to FLAIR (Figure 5).

**Figure 5.**
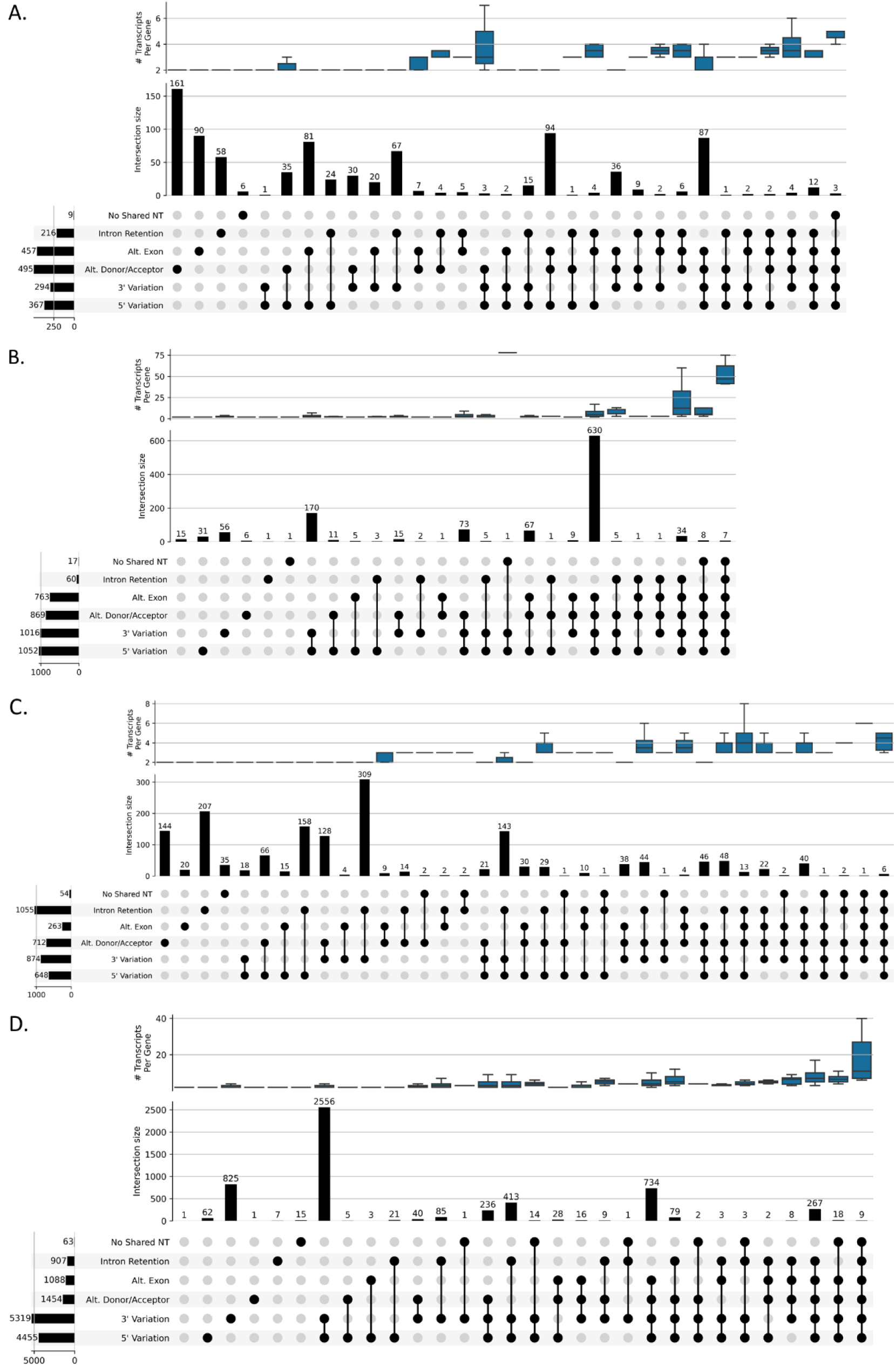
The number of genes with the specified types of alternative splicing indicated by the black dots below the histogram of gene counts for (A) *C. elegans* L1 FLAIR estimated transcriptome (n = 872 multi-transcript genes), (B) *C. elegans* L1 IsoSeq3 estimated transcriptome (n = 1159 multi-transcript genes), (C) *Z. mays* B73 root FLAIR estimated transcriptome (n = 1636 multi-transcript genes), and (D) *Z. mays* B73 root IsoSeq3 estimated transcriptome (n = 5464 multi-transcript genes). The box plots represent the number of transcripts per gene (blue). Genes with “No Shared NT” have a pair of transcripts with nonoverlapping coordinates.

We compared the estimated transcripts directly using 2GTF. FLAIR identified intron retention in *Z. Mays* and alterative exon cassettes/alternative donor acceptors in *C. elegans*, data respectively, as the primary mechanisms of splicing, recapitulating results from the species annotation comparisons in *TranD.* While IsoSeq3 identified 5’ and 3’ variation more frequently in both species. *TranD* (Figure 6, Supplementary Figure 8) We found that FLAIR is less likely to report small differences in splice sites, less likely to identify ambiguous junctions as different splice sites, and less likely to estimate transcripts that are an un-annotated subset of an existing longer transcript (Figure 7, Supplemental Figure 9) and IsoSeq3 is more likely to recognize a potentially novel transcript that is a subset of known exon combinations and more likely to identify multiple similar transcripts.

**Figure 6.**
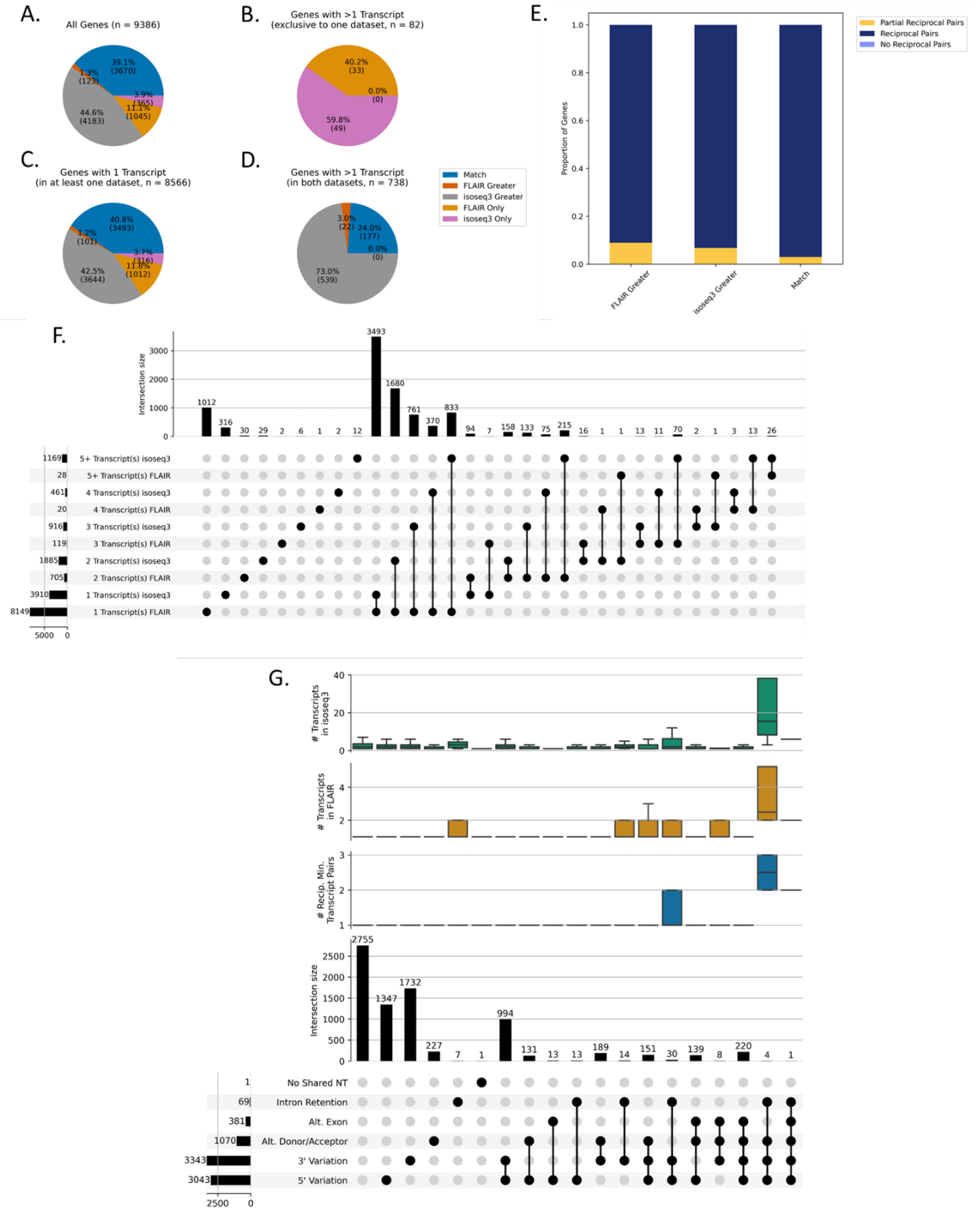
Transcripts are compared between *C. elegans* first larval stage (L1) and adult. (A-D) Percent of genes with equal number of transcripts between *C. elegans* L1 and adult (Match, blue), more transcripts in L1 compared to adult (L1 Greater, dark orange), more transcripts in adult compared to L1 (Adult Greater, gray), and genes exclusive to L1 (L1 Only, light orange) or adult (Adult Only, pink). (E) Within genes with equal numbers of transcripts in L1 and adult (Match), the proportion of genes where all transcripts have reciprocal minimum matches (Match: Reciprocal Pairs, n = 5896), at least one but not all transcript pairs are reciprocal minimum matches (Match: Partial Reciprocal Pairs, n = 70), and no pairs are reciprocal minimum matches (Match: No Reciprocal Pairs, n = 0). Within genes with more transcripts in L1 compared to adult (L1 Greater), the proportion of genes where all transcripts in adult have reciprocal minimum matches to a subset of L1 (L1 Greater: Reciprocal Pairs, n = 368), at least one but not all transcript pairs are reciprocal minimum matches (L1 Greater: Partial Reciprocal Pairs, n = 15), and no pairs are reciprocal minimum matches (L1 Greater: No Reciprocal Pairs, n = 0). Within genes with more transcripts in adult compared to L1 (Adult Greater), the proportion of genes where all transcripts in L1 have reciprocal minimum matches to a subset of adult (Adult Greater: Reciprocal Pairs, n = 520), at least one but not all transcript pairs are reciprocal minimum matches (Adult Greater: Partial Reciprocal Pairs, n = 23), and no pairs are reciprocal minimum matches (Adult Greater: No Reciprocal Pairs, n = 0). (F) Number of genes with the specified number of transcripts in L1 and adult indicated by the black dots below the histogram of genes counts. Columns with a single black dot represent the genes exclusive to L1 (n = 2129) or adult (n = 1738). Genes with more than one dot are in both L1 and adult (n = 6892). (G) Number of genes with the specified types of alternative splicing in only reciprocal minimum pairs between *C. elegans* first larval stage (L1) and adult indicated by the black dots below the histogram of gene counts (n = 6892 genes with 7304 reciprocal minimum pairs). Box plots represent the number of reciprocal minimum pairs (blue), number of transcripts in L1 (orange) and adult (green), and the average number (brown) and proportion (purple) of nucleotides different between the pairs. Genes with “No Shared NT” have a pair of transcripts with nonoverlapping coordinates.

**Figure 7.**
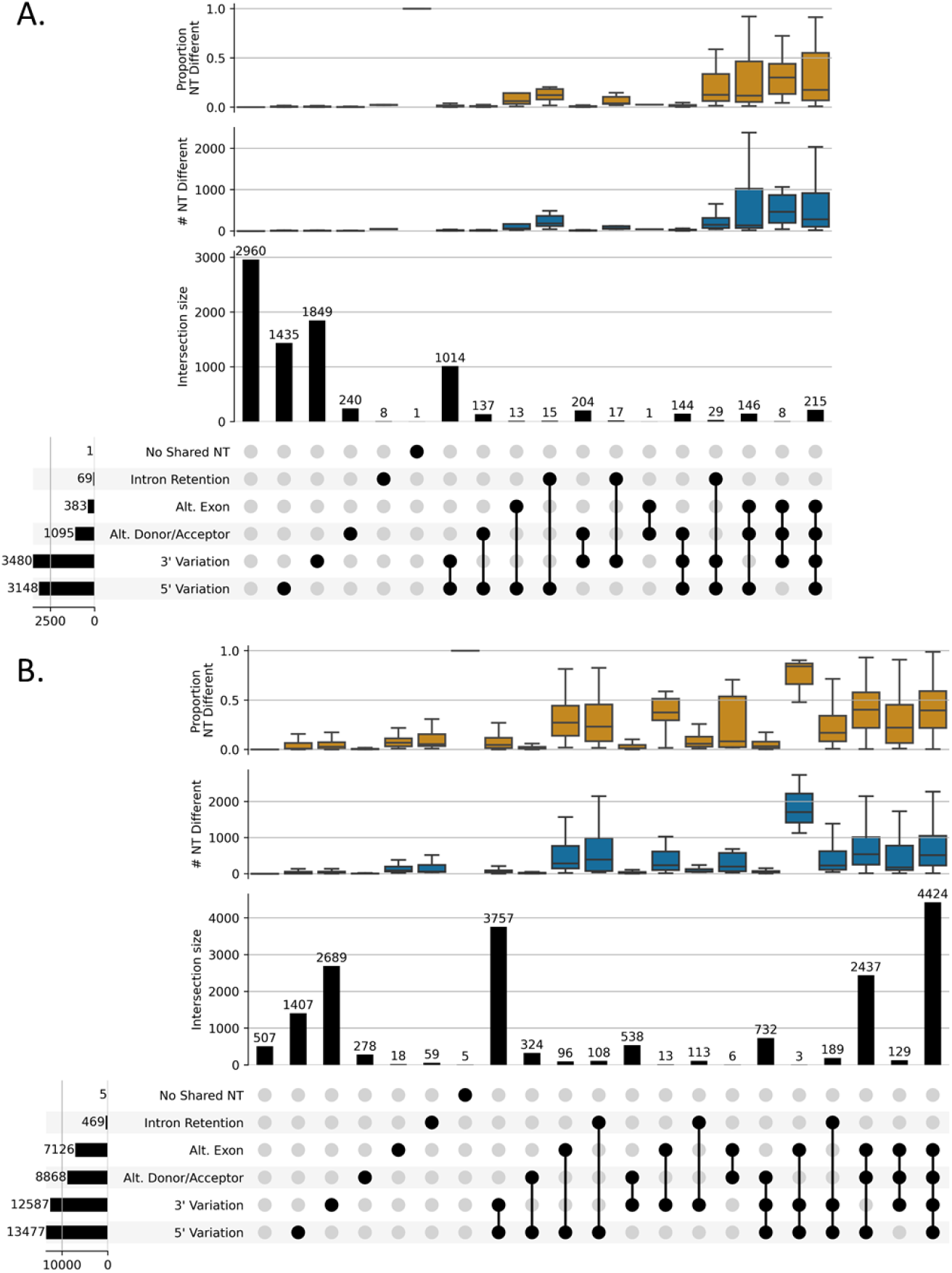
For *C. elegans* L1, the number of reciprocal minimum pairs (A, n = 8436 total pairs) and number of extra minimum pairs, or pairs without reciprocal minimums, (B, n = 17832 total pairs) with the specified types of alternative splicing between FLAIR and IsoSeq3 indicated by the black dots below the histogram of pair counts. Pairs with no shared nucleotides are in the same genes but with nonoverlapping coordinates. Box plots of the number (blue) and proportion (orange) of nucleotide (NT) differences between the pairs represented in the histogram.

Moreover, our analyses revealed that IsoSeq3 requires more reads to estimate transcripts and will filter out genes with few supporting reads (file *\*.cluster_report.csv* in the IsoSeq3 cluster output) (Figure 8) while FLAIR retains transcripts in genes if they are annotated as long as there are reads supporting that transcript. For researchers evaluating presence or absence of transcripts, and novel gene formation, such filtering may influence conclusions in evolutionary and biomedical applications. Both algorithms estimated an order of magnitude fewer transcripts than uniquely splicing reads.

**Figure 8.**
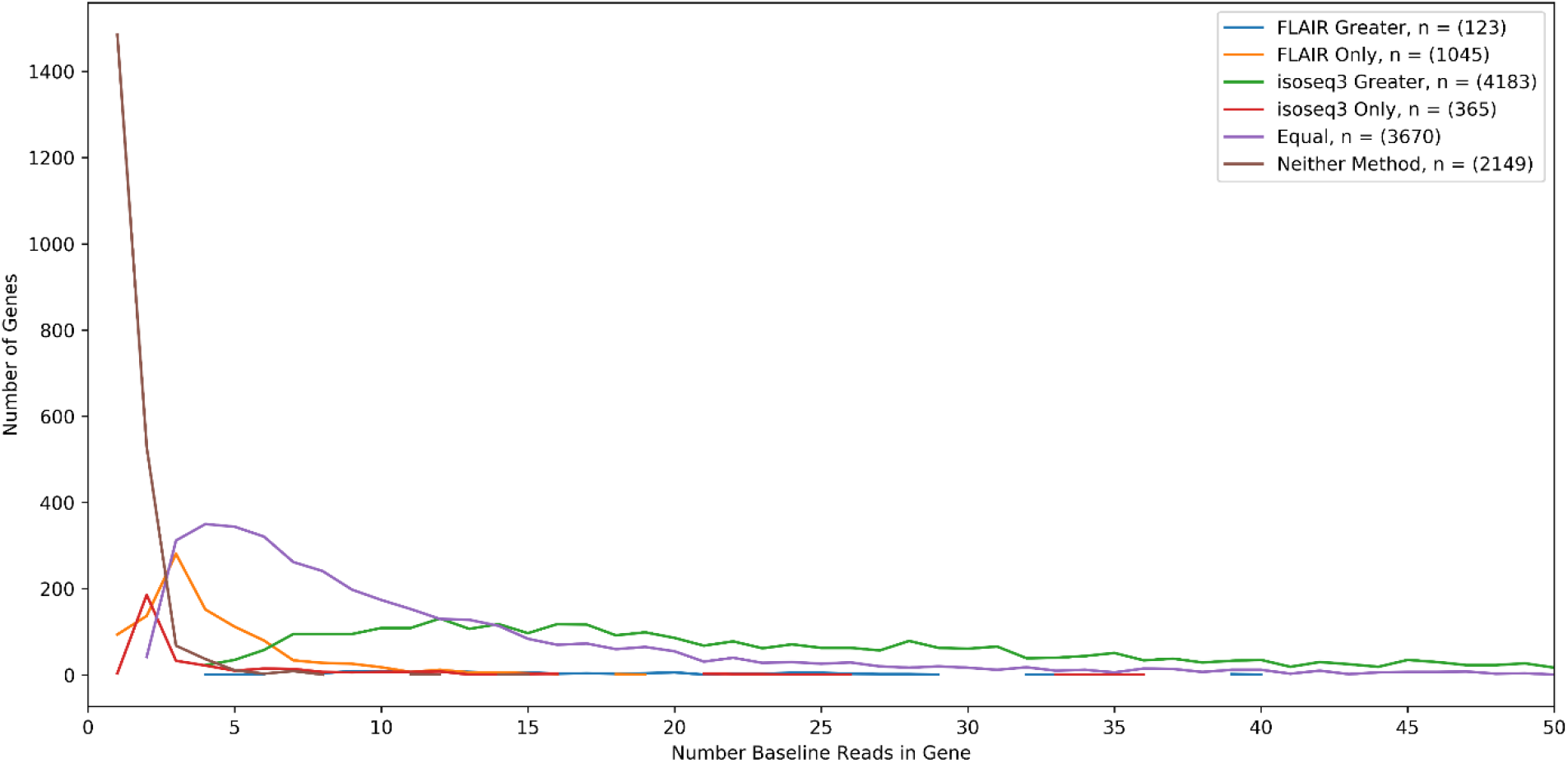
For *C. elegans* L1, the number of genes that contain the specified number of baseline reads (or selected consensus reads prior to transcript estimation) for each gene category determined by the number of transcripts estimated by FLAIR and IsoSeq3. A gene is categorized as “Neither Method” if a gene locus was detected by SQANTI3 QC in the baseline reads, but neither FLAIR nor IsoSeq3 estimated a transcript for the locus.

## Discussion

While there are a plethora of tools available to estimate (reviewed in Amarasinghe, et al. 2020) and consolidate/compare transcriptomes (Trapnell, et al. 2012; Kuo, et al. 2020) what is missing is a diagnostic tool that compares annotations and enumerates structural phenotypic differences between competing annotations. None of the existing tools are specifically designed to enumerate structural variation between transcripts. *TranD* enumerates structural variation between transcripts and pinpoints the nucleotide differences. It can describe transcripts within genes for a single GTF as well as enumerate the differences between all pairs of transcripts for the gene. In 2GTF mode all pairwise transcripts in 2 annotation files are compared for the same gene.

Annotation quality and completeness will always confound such comparisons. *TranD* is not a tool designed for quality control as these tools exist and are well established (Simao, et al. 2015; Tardaguila, et al. 2018). While this is a profoundly naïve assumption, and almost certainly violated in almost all cases, this particular limitation is not unique to this tool and is beyond the scope of this work. *TranD* does not do the hard work of identifying synteny and assigning common gene names (Haas, et al. 2003; Lyons and Freeling 2008) but rather assumes the gene names are already matched. The utility of *TranD* is to enumerate structural variation for the investigator. With few exceptions (Trapnell, et al. 2012; Tardaguila, et al. 2018), most approaches to describing similarity between transcripts rely on a version of BLAST (Altschul, et al. 1990) and often sophisticated (Yang, et al. 2018) functions of length comparisons that while they may account for variation in splicing patterns, the results do not necessarily list the nucleotide differences as a function of structural phenotypes. SQANTI (Tardaguila, et al. 2018) and Cuffcompare (Trapnell, et al. 2012) can be used to identify splice matches between a reference annotation and a new transcriptome model however SQANTI does not identify distances and Cuffcompare is primarily designed to track transcript fragments across multiple RNA-sequencing expression studies.

The map based approach that *TranD* uses is not designed to solve all known problems of sequence alignment: ambiguous junctions (Dehghannasiri, et al. 2019), map bias, and underlying inaccuracies in reference genomes are limitations shared by many bioinformatic pipelines (e.g., ExTraMapper Chakraborty, et al. 2021). Additional bioinformatic tools will be required to infer function, synteny, or phylogeny. Extramapper differs from *TranD* in that it deploys an algorithm with the goal of identifying a one-to-one orthologous transcript (or exon) between similar species, while *TranD* will compute all pairwise differences and calculates a minimum distance. TranD’s minimum distance is computed ‘with replacement’ such that more than one transcript in one GTF may map minimally to the same transcript in the second GTF. In addition, the metrics determining minimum distance are enumerated and returned as output along with the ability to calculate distances for all possible pairs, making it straightforward to change the minimum distance function.

Quantifying transcript differences begins with enumerating structural phenotypes: intron retention, alternative donors/acceptors, alternative exon cassettes, intron retentions, and alternative 3’ and 5’ ends. For each structural phenotype the number of shared nucleotides and variation in nucleotides is used to quantitate differences. This approach relies on being able to compare transcripts on the same coordinate system. In addition to distance metrics based on a single coordinate system, we develop metrics that describe transcriptome complexity that do not require identifying orthologs. We demonstrate that complexity metrics are effective in addressing fundamental questions relating the role of AS in evolution.

The advent of long reads has rapidly ushered in demand for the ability to understand, annotate and visualize long read data within and between species. Some important limitations to keep in mind when estimating transcriptomes from long read data: 1) the data are from a sample and so sampling variation should be expected, 2) there may be incompletely and imperfectly spliced molecules in the sample, 3) there may be ambiguous junctions that appear as alternate donor and acceptor sites (Dehghannasiri, et al. 2019), 4) there may be sequencing error, and 5) complexity of transcriptomes may be much higher than most annotations currently reflect. If the human annotation is correct, then perhaps other species should have more complexity in the number of transcripts than current models suggest. The excess variation in long reads may reflect true underlying complexity that has not yet been documented. Metrics that calculate the difference in transcripts to the nucleotide level enable individual researchers to assess the output of different modeling approaches. Complexity and distance metrics enable the researcher to compare estimates of the transcriptome to references, or to each other.

Cross-species comparisons under recent shared ancestry may be simple when homology allows for one-to-one matching. However, as species diverge in structure and in gene content, information from highly variable transcripts and newly emerged genes will be lost from analysis without clear orthologs. Total length and percent of shared sequence are not always the best methods for identifying a phylogenetic relationship. By comparing all pairs of transcripts and reporting the structural phenotypic differences between them when mapped to a common set of coordinates investigators can choose to weight certain types of structural phenotypes differently in choosing potential orthologs.

Genetic novelty is revealed by measuring distance between transcripts. Changes in transcriptomes between species offer a key source of evolutionary innovation. New exons that are formed through novel alternative splicing patterns or through acquisition of new genes may contribute to organism complexity, adaptive variation, or genetic disease (Bonnal, et al. 2020). Many modes of genetic innovation may reshape transcriptome content (polycistronic fusion genes, de novo genes, duplicate genes, chimeric genes, etc.), and rigorous methods are needed to ascertain their full effects at the molecular level. Better characterization of novel exon formation will help discern how these different sources of new genetic structure in the transcriptome may contribute to neutral, adaptive, and detrimental variation. *TranD* will identify these structural phenotypes and identify them relative to other transcripts in all genes in the GTF file.

## Acknowledgements

R01GM128193 (LMM), Srna Vlaho, Sarah Signor, Sergey Nuzhdin (fly collections), the department of Molecular Genetics and Microbiology (sequencing costs), University of Florida Research Computing, HiPerGator, MIRA R35 GM133376 (RLR), Jeremy R.B. Newman for questions and event analysis, Alison Morse for testing, user guide and comments

## Notes

https://github.com/McIntyre-Lab/TranD

### Competing Interest Statement

The authors have declared no competing interest.

